# Disruption of Hepatic Mitochondrial Pyruvate and Amino Acid Metabolism Impairs Gluconeogenesis and Endurance Exercise Capacity in Mice

**DOI:** 10.1101/2023.08.22.554345

**Authors:** Michael R. Martino, Mohammad Habibi, Daniel Ferguson, Rita T. Brookheart, John P. Thyfault, Gretchen A. Meyer, Louise Lantier, Curtis C. Hughey, Brian N. Finck

## Abstract

Exercise robustly increases the glucose demands of skeletal muscle. This demand is met not only by muscle glycogenolysis, but also by accelerated liver glucose production from hepatic glycogenolysis and gluconeogenesis to fuel mechanical work and prevent hypoglycemia during exercise. Hepatic gluconeogenesis during exercise is dependent on highly coordinated responses within and between muscle and liver. Specifically, exercise increases the rate at which gluconeogenic precursors such as pyruvate/lactate or amino acids are delivered from muscle to the liver, extracted by the liver, and channeled into glucose. Herein, we examined the effects of interrupting gluconeogenic efficiency and capacity on exercise performance by deleting hepatic mitochondrial pyruvate carrier 2 (MPC2) and/or alanine transaminase 2 (ALT2) in mice. We found that deletion of MPC2 or ALT2 alone did not significantly affect time to exhaustion or post-exercise glucose concentrations in treadmill exercise tests, but mice lacking both MPC2 and ALT2 in liver (DKO) reached exhaustion faster and exhibited lower circulating glucose during and after exercise. Use of ²H/¹³C metabolic flux analyses demonstrated that DKO mice exhibited lower endogenous glucose production owing to decreased glycogenolysis and gluconeogenesis at rest and during exercise. The decreased gluconeogenesis was accompanied by lower anaplerotic, cataplerotic, and TCA cycle fluxes. Collectively, these findings demonstrate that the transition of the liver to the gluconeogenic mode is critical for preventing hypoglycemia and sustaining performance during exercise. The results also illustrate the need for interorgan crosstalk during exercise as described by the Cahill and Cori cycles.

## INTRODUCTION

Exercise evokes dramatic changes in skeletal muscle and systemic metabolism that allow the organism to meet the energy demands of muscular work. Muscle glycogen serves to fuel muscle contraction, but muscle must also rely on blood glucose and other metabolic substrates to fuel contractions. The liver is vital to the maintenance of adequate blood glucose concentrations during exercise and for replenishing muscle glycogen after exercise under fasting conditions (20, 30). Hepatic glucose output is the product of hepatic glycogenolysis and gluconeogenesis. Gluconeogenesis, the synthesis of new glucose from non-carbohydrate precursors (lactate/pyruvate, glycerol, and gluconeogenic amino acids), becomes increasingly important as liver glycogen levels are expended during prolonged exercise (5, 27, 30). Indeed, liver-specific deletion of cytosolic phosphoenolpyruvate carboxykinase (PEPCK1), which catalyzes a rate limiting step in gluconeogenesis, leads to impaired exercise performance in mice (21).

The stimulation of liver gluconeogenesis by exercise is dependent on highly coordinated metabolic responses within and between muscle and liver. It requires an increase in the rate at which gluconeogenic precursors are delivered from muscle to the liver, extracted by the liver, and channeled into glucose. Work by Gertrude and Carl Cori defined interorgan metabolic crosstalk, which now bears their name (the Cori cycle), that is foundational for glucose metabolism under multiple conditions including exercise (22). During exercise, skeletal muscle anaerobic glucose metabolism and the production of pyruvate/lactate are elevated. Much of this lactate enters the bloodstream and is taken up by the liver for use as a gluconeogenic substrate. Glucose produced from this process can then be released by the liver and then travel through the blood to skeletal muscle to be metabolized via glycolysis.

The Cahill cycle is analogous in nature to the Cori cycle, but involves shuttling alanine produced by peripheral tissues to the liver for its use in hepatic gluconeogenesis. Alanine release by muscle during exercise is increased (31) and alanine is present in blood far in excess of its proportional abundance to other amino acids in skeletal muscle (7, 8). Alanine can be generated from breakdown of liver and muscle protein (23), but transamination of amino groups to pyruvate in muscle is likely a primary source of alanine; especially in the context of physiologic stimuli like exercise (25, 26). Muscle-synthesized alanine is delivered to the liver (31) where the amino group is removed and funneled to the urea cycle while re-synthesizing pyruvate, which can then be converted to glucose. Ultimately, the Cahill Cycle (6) serves two important functions during exercise; it delivers gluconeogenic precursors and muscle-derived amino groups in the nontoxic form of alanine to the liver.

The Cori and Cahill cycles and their connection to hepatic gluconeogenesis have been well-studied. However, the regulatory nodes underlying their connection to hepatic gluconeogenesis have yet to be clearly defined. For example, the mitochondrial pyruvate carrier (MPC) is a heterodimeric inner mitochondrial membrane protein complex composed of two proteins (MPC1 and MPC2) that catalyzes the mitochondrial import of pyruvate (2, 11). This step is critical for pyruvate entry into the gluconeogenic pathway because pyruvate carboxylase, which catalyzes a required step in pyruvate mediated gluconeogenesis (3), is exclusively localized in the mitochondrial matrix. Mice lacking MPC proteins in hepatocytes are protected from hyperglycemia in diabetic mouse models due to reduced gluconeogenic flux (9, 16) and MPC inhibitors are also known to lower blood glucose in these models (4, 12). Similarly, alanine transaminase (ALT) enzymes are required for alanine to be converted to pyruvate and enter the gluconeogenic pathway via pyruvate carboxylase. Importantly, two distinct isozymes of ALT (ALT1 and ALT2) are encoded by two genes in *Mammalia* (*Gpt* and *Gpt2*). ALT1 is cytosolic while ALT2 is localized to the mitochondria (10, 14, 19). Recent work has demonstrated that hepatic ALT protein expression is increased in mice and humans with obesity and that suppressing the expression of ALT enzymes in liver has anti-diabetic effects by reducing alanine-stimulated gluconeogenesis (15, 18). Herein, we tested the effects of liver MPC and ALT2 deletion on hepatic nutrient metabolism during exercise and its impact on exercise performance. Studies used mice deficient in hepatic MPC2, ALT2, or both MPC2 and ALT2 (DKO) and evaluated their performance in exercise endurance and capacity tests. These indicators of exercise performance were accompanied by the quantification of *in vivo* glucose and associated nutrient fluxes at rest and during exercise.

## RESEARCH DESIGN AND METHODS

### Animal Studies

All the experiments were performed with 8- to 16-week-old mice of both sexes. At the conclusion of exercise experiments, mice were sacrificed by injection of 150 mg/kg body weight of sodium pentobarbital “Fatal Plus” (Vortex Pharmaceuticals, Dearborn MI) following the final blood glucose and lactate collection time point prior to tissue collection. For all other experiments, mice were sacrificed with CO_2_. All the experiments involving mice were approved by the Institutional Animal Care and Use Committees of Washington University in St. Louis and Vanderbilt University and are in agreement with the *Guide for the Care and Use of Laboratory Animals*.

### Generation of DKO mice

The generation LS-*Mpc2*-/- and LS-*Gpt2*-/- mice has been described (15, 16). *Gpt2* fl/fl mice were mated with LS-*Mpc2*-/- mice to generate LS-*Gpt2/Mpc2* -/- (DKO) mice (32). Littermate mice not expressing Cre (fl/fl mice or double *Mpc2*/*Gpt2* fl/fl mice) were used as control mice in all experiments.

### Exercise protocol (for studies shown in Figure 1, 2, and 4)

For exercise studies shown in Figures 1, 2, and 4, mice were run to exhaustion on a closed treadmill (Columbus Instruments) with a shock grid that delivered a mild electrical stimulus (16-28 V) to encourage continuous running. Food was removed 4.5 h before exercise, bedding was replaced with aspen chip bedding and initial body weights were recorded. Mice were weighed immediately before the run and then acclimated to the treadmill with 0° incline at 0 m/min for 5 min. The speed was then increased to 5 m/min and maintained for 5 min. After 5 min of continuous running speed was increased to 10 m/min, after 10 min speed was increased to 15 m/min, after 20 min speed was increase to 25 m/min and after 35 min the speed was increase to 30 m/min and maintained until the mice reached exhaustion. Exhaustion was determined by refusal of mice to remain on the treadmill belt for 10 seconds and lack of movement upon removal from the treadmill. Blood glucose and lactate were measured with a One-Touch Ultra glucometer (LifeScan) and a Lactate Plus lactate meter (NovaBiomedical), respectively, from a single drop of blood from the tail vein. Blood glucose and lactate concentrations were measured immediately upon exhaustion (T=0), and at T=5, 10, 15, 30, and 60 min post exhaustion. Mice remained fasted during the 1 h post-exercise period.

**Figure 1.**
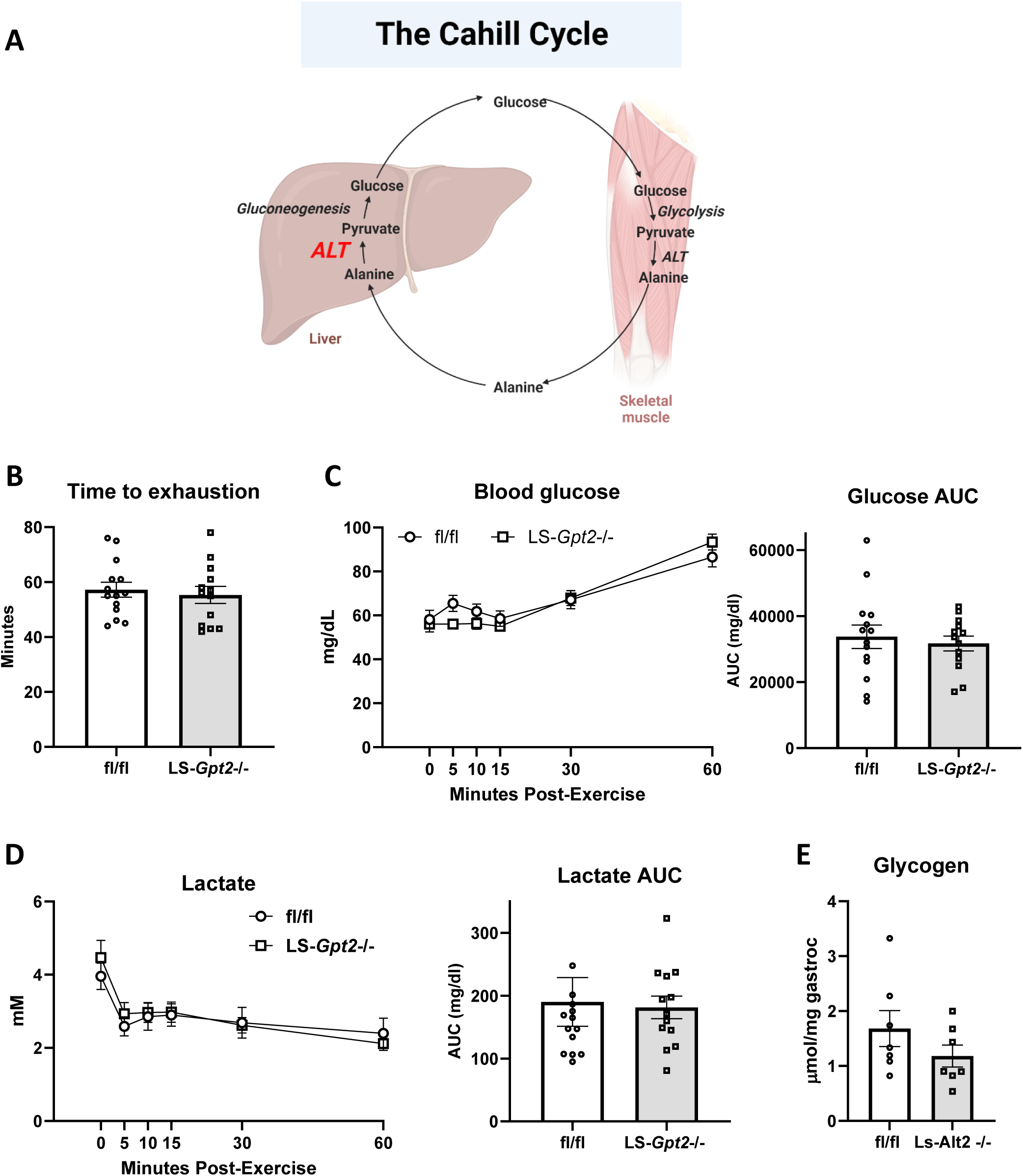
(A) Schematic depiction of the Cahill Cycle, wherein skeletal muscle derived alanine is converted to glucose in the liver. Created using BioRender.com. (B) Time taken to reach exhaustion during a graded exercise test in fl/fl and LS-*Gpt2-/-* mice. (C and D) Blood glucose (C) and blood lactate (D) concentrations and area under the curve (AUC) in fl/fl and LS-*Gpt2-/-* mice during the 60 min post-exercise recovery period. (E) Gastrocnemius glycogen in fl/fl and LS-*Gpt2-/-* mice at 60-min post-exercise.

**Figure 2.**
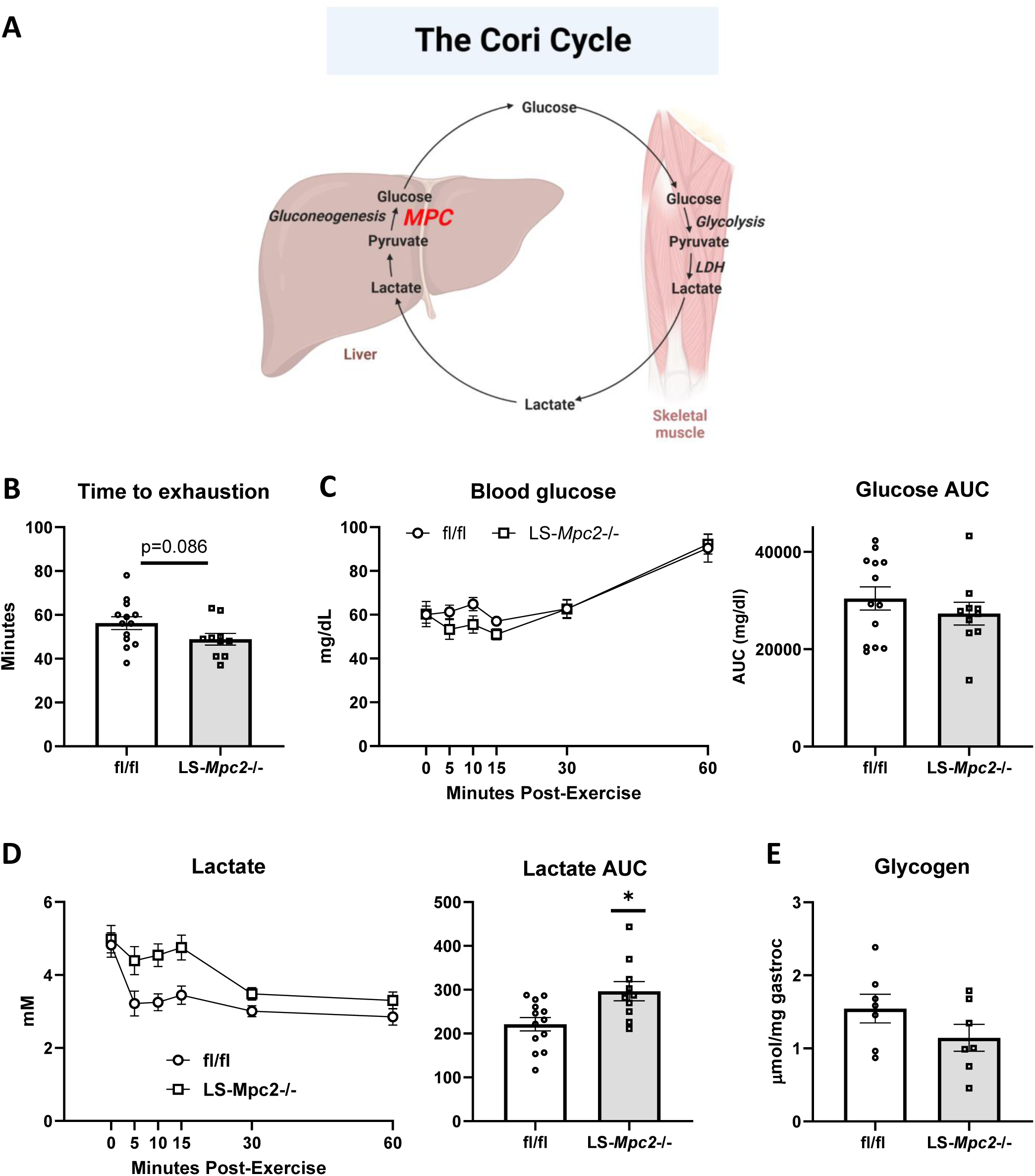
(A) Schematic depiction of the Cori Cycle, wherein skeletal muscle derived lactate is converted to glucose in the liver. Created using BioRender.com. (B) Time taken to reach exhaustion during a graded exercise test in fl/fl and LS-*Mpc2-/-* mice. (C and D) Blood glucose (C) and blood lactate (D) concentrations and area under the curve (AUC) in fl/fl and LS-*Mpc2-/-* mice during the 60 min post-exercise recovery period. (E) Gastrocnemius glycogen in fl/fl and LS-*Mpc2-/-* mice at 60-min post-exercise. Data are presented as mean ± SEM.

**Figure 3.**
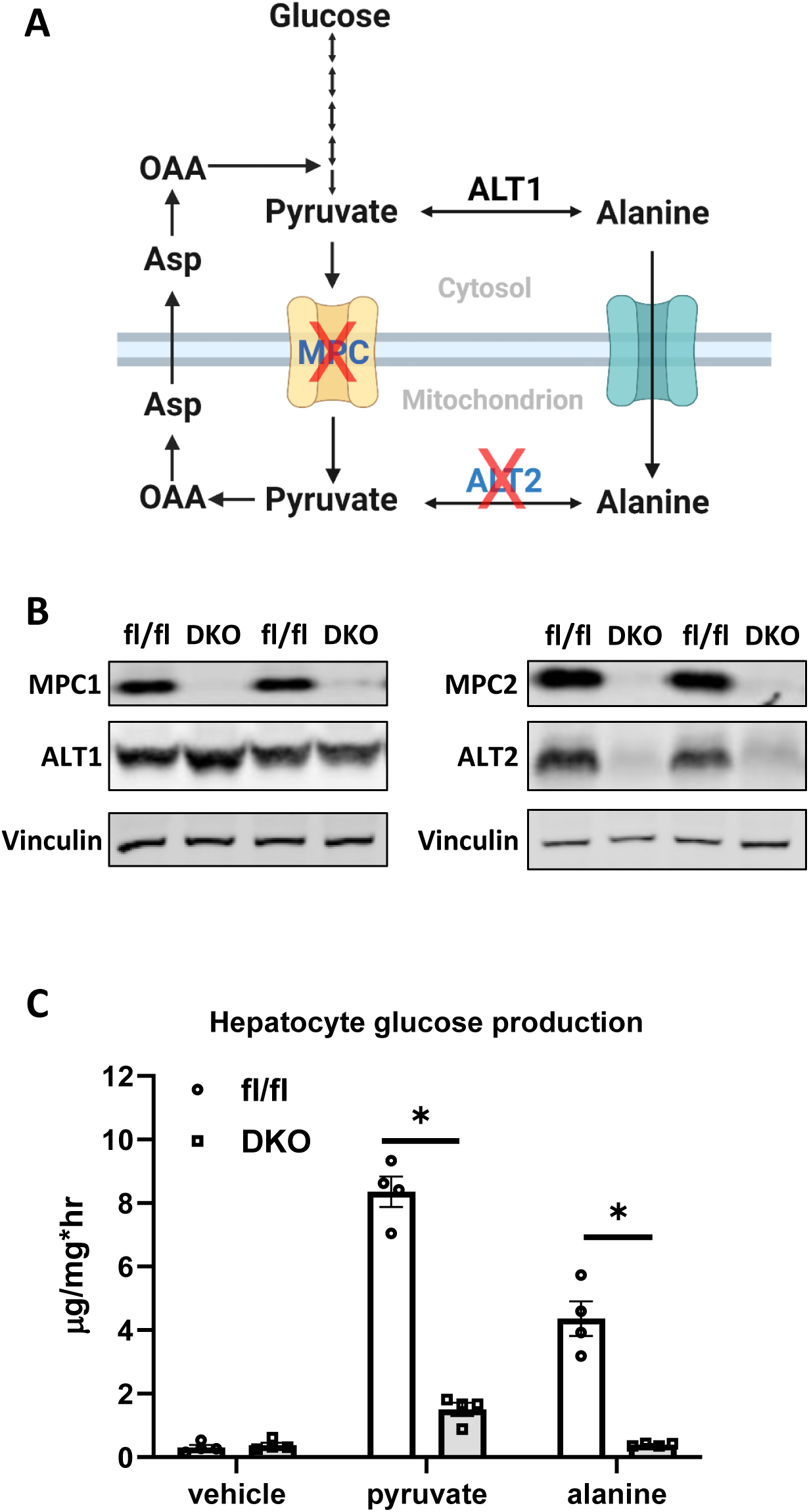
(A) Schematic representation of gluconeogenesis from pyruvate and alanine and the potential impact of deleting the MPC and ALT2 enzymes. Created using BioRender.com. (B) Representative Western blots using liver protein lysates from fl/fl and DKO mice using the indicated antibodies. (C) Glucose concentrations in the media of cultured hepatocytes stimulated with glucagon in the presence of gluconeogenic substrates. Pyr – pyruvate (5 mM), Ala – alanine (20 mM). Data are presented as mean ± SEM. *p<0.05 vs fl/fl control.

**Figure 4.**
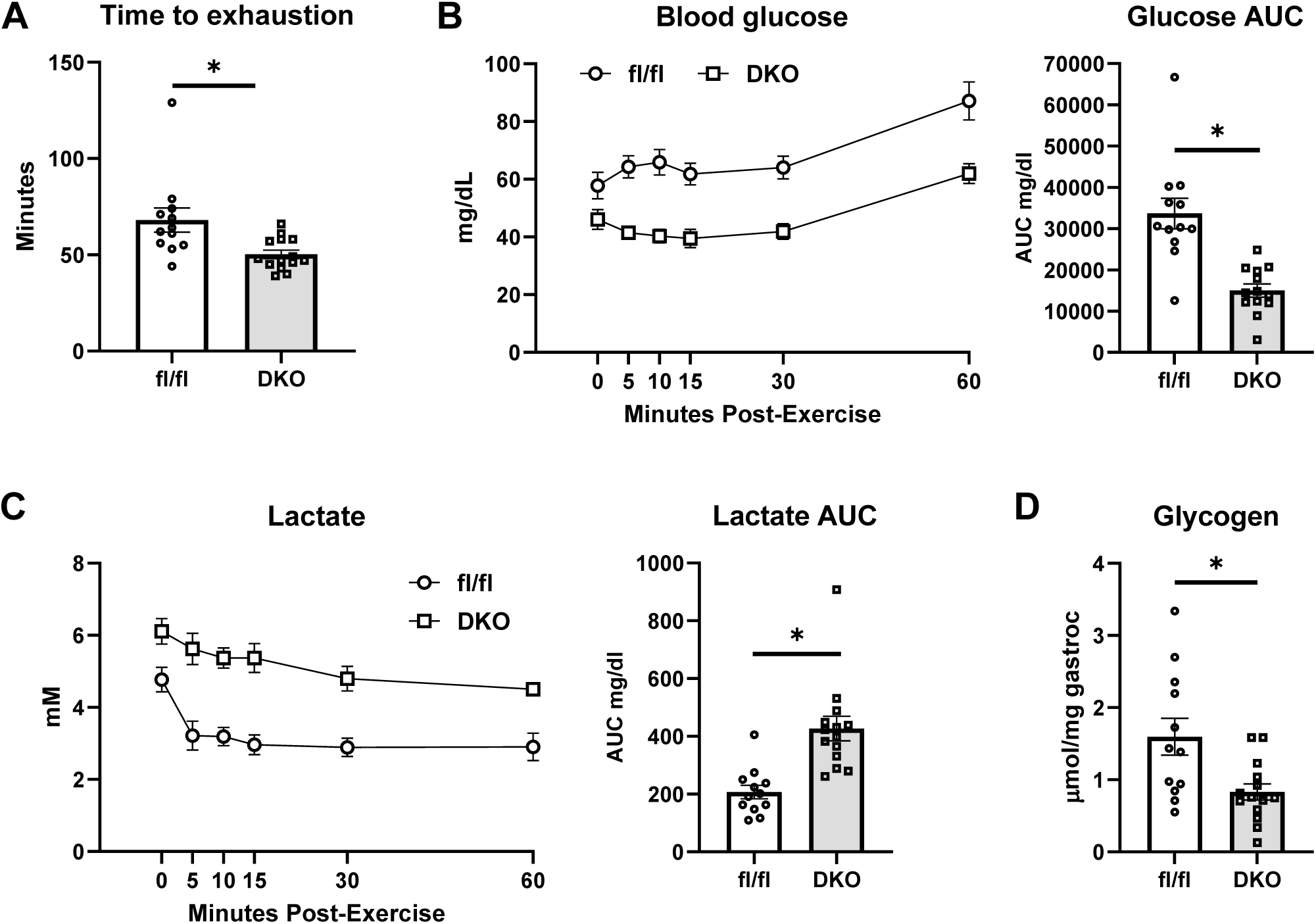
(A) Exercise time to exhaustion in fl/fl and DKO mice. (B and C) Blood glucose (B) and blood lactate (C) concentrations and area under the curve (AUC) in fl/fl and DKO mice during the 60 min post-exercise recovery period. (D) Glycogen concentrations of gastrocnemius muscles taken 60-min post exercise in fl/fl and DKO mice. Data are presented as mean ± SEM. *p<0.05.

### Exercise protocol (for studies shown in Figure 5 and 6)

Prior to studies shown in Figure 5 and 6, mice were transferred to the Vanderbilt University Mouse Metabolic Phenotyping Core. After 4 weeks of quarantine and acclimation, mice underwent an exercise stress test to assess maximal running speed. In this paradigm, the treadmill speed starts at 10 m/min and the speed increases 4 m/min every 3 min until exhaustion (Supplemental Figure 1). Thereafter, catheters were implanted in the jugular vein and carotid artery for stable isotope infusion and sampling protocols as previously described (1, 13, 21). The free ends of the catheters were tunneled under the skin to the back of the neck and the exteriorized ends of the vascular catheters were flushed with 200 U/ml of heparinized saline and sealed with stainless-steel plugs. Following the surgical procedures, mice were individually housed and provided ∼9-10 days of post-operative recovery prior to stable isotope infusions studies during rest and acute exercise.

**Figure 5.**
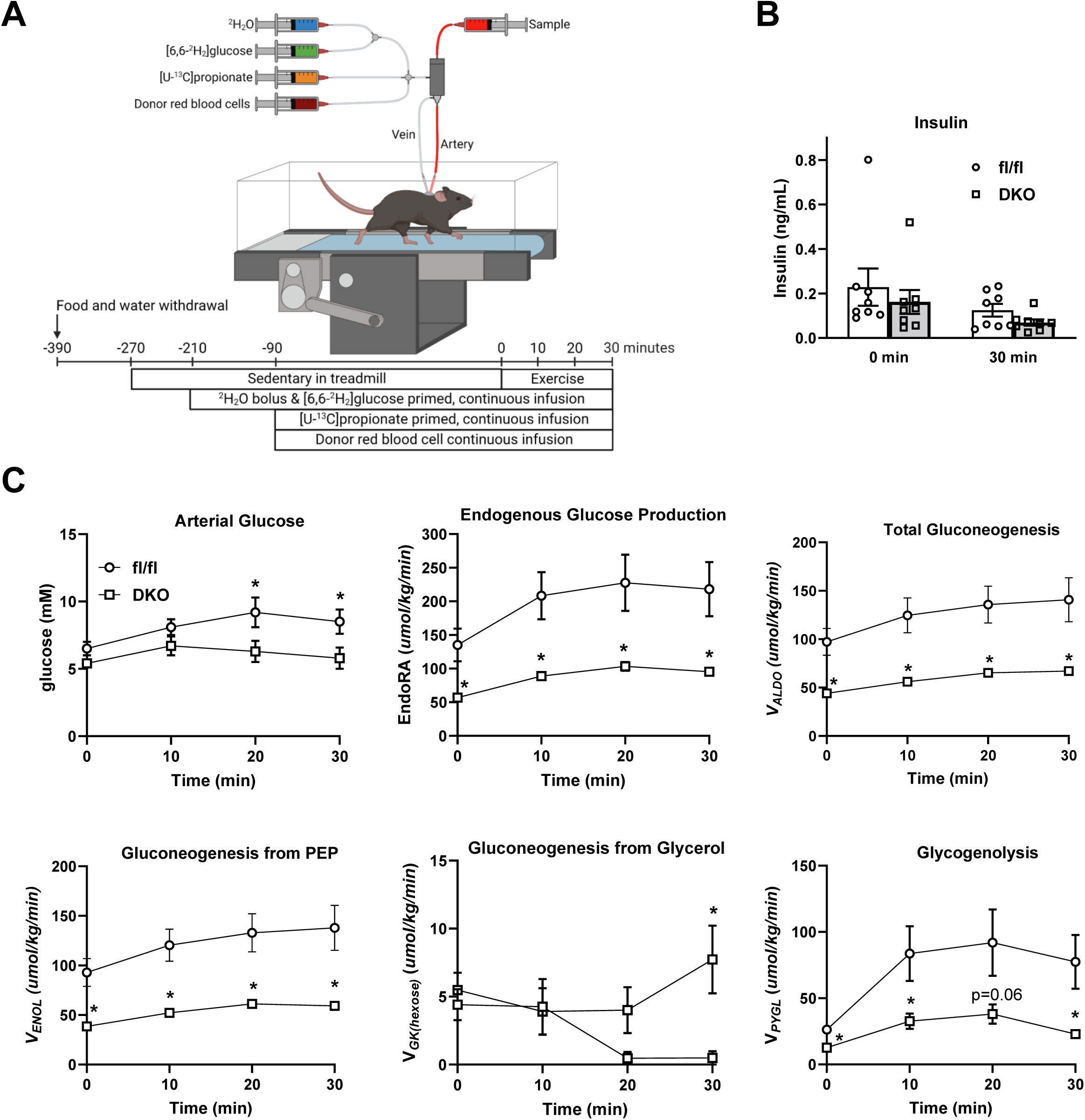
(A) Exercise experiment schematic. Created using BioRender.com. Stable isotope infusions at rest and during acute treadmill running bout were performed in mice ∼9-10 days following carotid arterial and jugular catheter implantation surgeries. At 210 min prior to the treadmill running bout (3 h of fasting), a ^2^H_2_O bolus was administered into the venous circulation to enrich total body water at 4.5%. A [6,6-^2^H_2_]glucose prime was infused followed by a continuous infusion was initiated with the ^2^H_2_O bolus. Ninety min before the onset of exercise (5 h of fasting), a primed, continuous infusion of [U-^13^C]propionate was started. Donor red blood cells were administered to prevent a decline in hematocrit. Arterial samples were obtained prior to stable isotope infusion as well as during 30 min exercise bout for ^2^H/^13^C metabolic flux analysis. (B) Plasma insulin concentrations at baseline (0 min) and after 30 min of exercise. (C) A time course of blood glucose concentration (mmol/L) in fl/fl and DKO mice prior to (0-min time point) and during a 30 min treadmill run (10-30 min time points). Model-estimated, nutrient fluxes (μmol·kg-1·min-1) in fl/fl and DKO mice prior to and during a 30-min of treadmill run for endogenous glucose production, total gluconeogenesis (VAldo), gluconeogenesis from phosphoenolpyruvate (VEnol), gluconeogenesis from glycerol (VGK), and glycogenolysis (VPYGL). *p<0.05 vs. fl/fl at specified time point by two-way repeated measures ANOVA followed by Šidák’s post-hoc tests.

**Figure 6.**
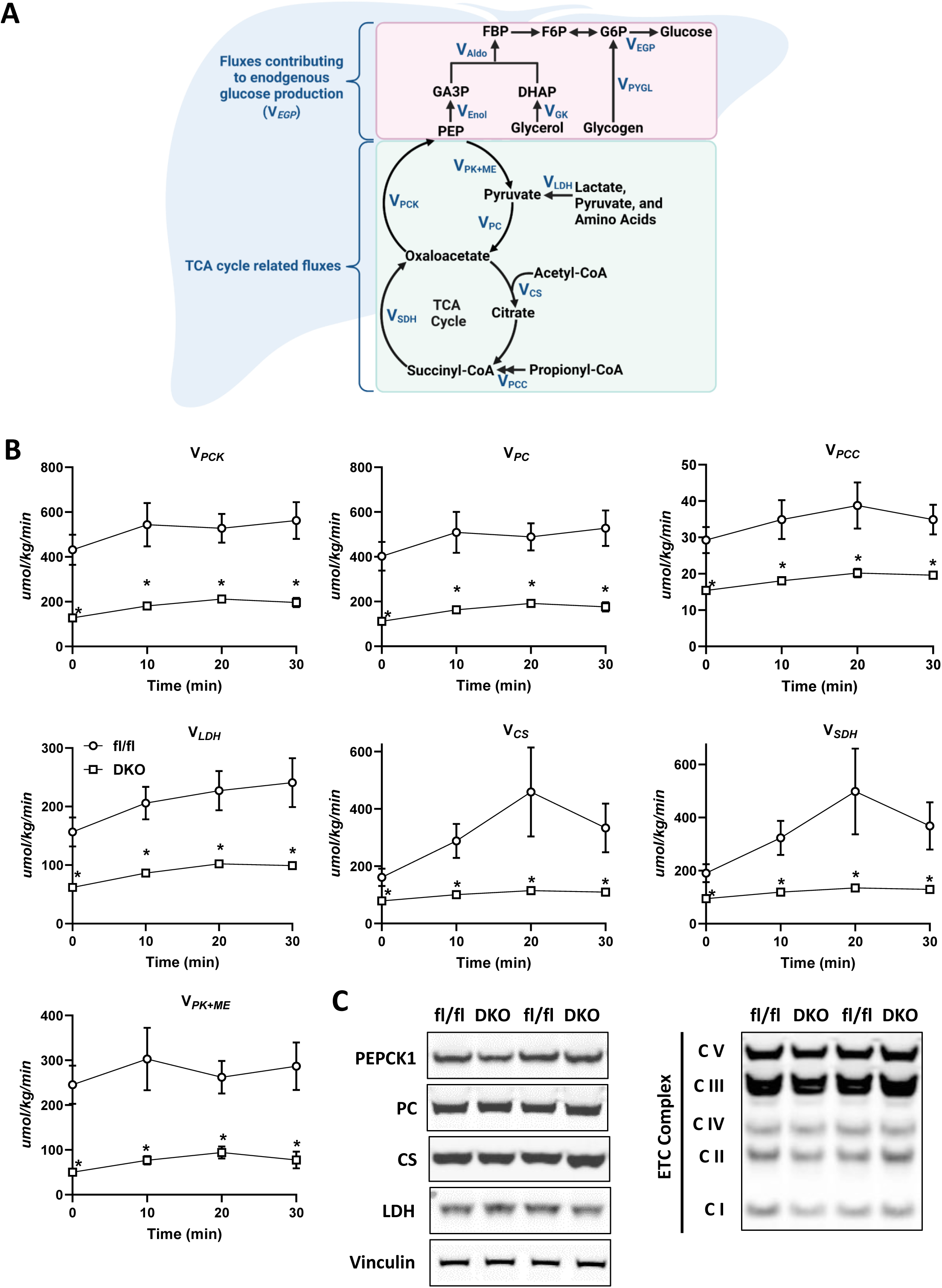
(A) Schematic representation of select glucose producing and tricarboxylic acid cycle fluxes quantified by ^2^H/^13^C metabolic flux analysis. Created using BioRender.com. (B) Model-estimated, nutrient fluxes (μmol·kg-1·min-1) in fl/fl and DKO mice prior to and during a 30-min of treadmill run for for total cataplerosis (VPCK), anaplerosis from pyruvate (VPC), anaplerosis from propionyl-CoA (VPCC), flux from unlabeled, non-phosphoenolpyruvate, anaplerotic sources to pyruvate (VLDH), flux from oxaloacetate and acetyl-CoA to citrate (VCS), flux from succinyl-CoA to oxaloacetate (VSDH), and pyruvate cycling (VPK+ME). Data are mean ± SEM. *p<0.05 vs. fl/fl at specified time point by two-way repeated measures ANOVA followed by Šidák’s post-hoc tests. (C) Representative Western blots using liver lysates from fl/fl and DKO mice after completing this exercise protocol using the indicated antibodies.

The weights of all mice were within 10% of pre-surgery body weight prior to the stable isotope infusions and the exercise protocol. Food and water were withdrawn within 1 h of start of light cycle (6-8 am). Two h into the fast, mice were placed in an enclosed single lane treadmill (Columbus Instruments, Columbus, OH) and the exteriorized catheters were connected to infusion syringes (Figure 4). Three h into the fast, an 80 μl arterial blood sample was obtained to determine natural isotopic enrichment of plasma glucose. Immediately following this sample, stable isotope infusions were initiated as previously performed (13, 21) (Figure 5). Briefly, a ^2^H_2-_ O (99.9%)-saline bolus containing [6,6-^2^H_2_]glucose (99%) was administered over 25 min to both enrich body water and deliver a [6,6-^2^H_2_]glucose prime (440 μmol/kg). This was immediately followed by a continuous infusion of [6,6-^2^H_2_]glucose (4.4 μmol/kg/min). A primed (1.1 mmol/kg), continuous (0.055 mmol/kg/min) intravenous infusion of [U-^13^C]propionate was started two h after the ^2^H_2_O bolus and [6,6-^2^H_2_]glucose prime. Four 100-150 μl arterial blood samples were obtained (90-120 min following the [^13^C_3_] propionate bolus) to determine arterial blood glucose concentration as well as plasma glucose enrichment used in ^2^H/^13^C metabolic flux analyses protocols to quantify hepatic glucose and associated nutrient fluxes. The sample taken at 90 min following the [U-^13^C]propionate bolus (Time = 0 min) was obtained while mice were at rest on a stationary treadmill. Samples taken 100-120 min following the [U-^13^C]propionate bolus (Time = 10-30 min) were obtained while mice were performing a 30-min acute treadmill running bout at 12 m/min. Donor red blood cells were given by constant rate infusion for the duration of the study to ensure hematocrit did not fall more than 10%. Immediately following the exercise bout, mice were sacrificed via pentobarbital and tissue collected was flash frozen.

### Glucose Derivatizations and GC-MS

Forty μl of plasma from the -210-, 0-, 10-, 20-, and 30-min time points was processed to obtain di-*O*-isopropylidene propionate, aldonitrile pentapropionate, and methyloxime pentapropionate derivatives of glucose as previously described (13). Briefly, GC-MS injection volumes of 1 μl with purge flow times between 20 and 120 seconds was used. A custom MATLAB function was used to integrate derivative peaks in order to obtain mass isotopomer distributions (MIDs) for six glucose fragment ions. The following fragment ion ranges were used for determining uncorrected MIDs: aldonitrile, m/z 173–178, 259–266, 284–291, and 370–379; methyloxime, m/z 145–149; di-O-isopropylidene, m/z 301–314.

### ^2^H/^13^C Metabolic Flux Analysis

The *^2^H/^13^C Metabolic Flux Analysis* methodology has been previously described (13). To summarize, a reaction network was generated using Isotopomer Network Compartmental Analysis (INCA) software. The reaction network defined carbon and hydrogen transitions for hepatic glucose producing and associated oxidative metabolism reactions. Flux through each network reaction was determined relative to citrate synthase flux (V_CS_) by minimizing the sum of squared residuals between simulated and experimentally determined MIDs of the six fragment ions. Flux estimates were repeated 50 times from random initial values, goodness of fit was evaluated by a chi-square test (*p* = 0.0*5*), and confidence intervals of 95% were determined. Mouse body weights and the [6,6-^2^H_2_]glucose infusion rate were used to quantify absolute flux values.

### Muscle glycogen quantification

Skeletal muscle tissue was collected at sacrifice and snap frozen in liquid nitrogen. Glycogen concentration was determined using frozen tissues (30 to 90 mg) hydrolyzed in 0.3 ml of 30% (wt/vol) KOH solution in a boiling water bath for 30 min. At 10 and 20 min of the incubation, tubes were shaken by hand to facilitate the digestion. After cooling to room temperature, 0.1 ml of 1 M Na_2_SO_4_ and 0.8 ml of ethanol were added, the samples were boiled again for 5 min to facilitate precipitation of glycogen and then centrifuged at 10,000 x *g* for 5 min. The glycogen pellet was dissolved in 0.2 ml of water, and two additional ethanol precipitations were performed. The final pellet was dried and dissolved in 0.2 ml of 0.3-mg/ml amyloglucosidase in 0.2 M sodium acetate buffer (pH 4.8) and incubated for 3 h at 40°C. The reaction mixture was diluted two- to fivefold with water. To determine the glucose concentration, 5 ml of the diluted sample was added to 0.2 ml of the glucose assay solution which contains 0.3 M triethanolamine–KOH (pH 7.5), 1 mM ATP, 0.9 mM b-NADP, and 5 mg of G-6P dehydrogenase per ml. The absorbance at 340 nm was determined before and after addition of 1 mg of hexokinase. Glycogen content is expressed as micromoles of glucosyl units per gram (wet weight) (28).

### Protein isolation and western blotting

For western blotting, liver lysates were collected in RIPA lysis buffer (Cell Signaling Technology; 9806) with protease/phosphatase inhibitors (Cell Signaling Technology; 5872) using a TissueLyser. Lysates were normalized to protein concentration, denatured, and run on NuPAGE precast 4-12%gels (ThermoFisher Scientific) using MOPS or MES buffers (ThermoFisher Scientific). Separated proteins were transferred to Immobilon PVDF membrane and blocked for one hour with 5% BSA in TBST. The antibodies used in this study were ALT1 (Abcam; ab154034, ALT2 (Abcam; ab202083), MPC1 (Cell Signaling Technology; D2L9I), MPC2 (Cell Signaling Technology; D4I7G), Vinculin (Cell Signaling Technology; 13901), PEPCK (Cayman Chemical; 10004943), PC (Cell Signaling Technology; 66470), CS (Cell Signaling Technology; 14309), LDH (Abcam; ab52488), OXPHOS cocktail (Abcam; ab110413). Blots were imaged using a LI-COR Odyssey system with Image Studio Lite software.

### Hepatocyte Isolation

Hepatocytes were isolated by perfusing livers of anesthetized mice with DMEM media containing collagenase from *Clostridium histolyticum* (Sigma Chemical Co.) as reported before (16). Briefly, hepatocytes were plated overnight in collagen coated 12 well-plates (200,000 cells/mL) with DMEM containing 10% FBS plus an antibiotic cocktail (penicillin/streptomycin and amphotericin) and washed twice the following morning with glucose-free Hank’s balanced salt solution (GF-HBSS) containing 127mM NaCl, 3.5 mM KCl, 0.44 mM KH_2_PO_4_, 4.2 mM NaHCO_3_, 0.33 mM Na_2_HPO_4_, 1 mM CaCl_2_, 20 mM HEPES, pH 7.4.

### Glucose production assay

Following the final wash, hepatocytes were starved for 2 h in GF-HBSS. HBSS was removed, cells were washed in fresh HBSS, and then treated for 3 h in HBSS containing glucagon (100 ng/mL) alone or 5 mM sodium pyruvate or 20 mM alanine. After the 3 h incubation, media was collected and glucose concentrations were measured using a glucose oxidase-based glucose assay kit (MilliporeSigma; cat# GAGO20). Glucose concentrations were normalized to cell protein amount measured by Micro BCA kit (ThermoFisher).

### Statistical Analyses

Figures were prepared using Prism version 8.0.1 for Windows (GraphPad Software, La Jolla California USA, www.graphpad.com). All data are presented as the mean ± SE. Statistical significance was calculated using an unpaired Student’s *t*-test or two-way analysis of variance (ANOVA) with repeated measures for data analysis, with a statistically significant difference defined as *p* ≤ 0.05.

## RESULTS

### Loss of liver ALT2 or MPC2 alone does not limit exercise performance

The conversion of alanine to glucose by the liver is believed to be important to maintaining blood glucose concentrations during and after exercise when food is not available via the Cahill Cycle (Figure 1A). We recently generated mice with liver-specific deletion of *Gpt2*, which encodes the mitochondrial alanine transaminase ALT2, and have reported their phenotype at baseline and after prolonged fast (15). Briefly, these mice are overtly normal and did not exhibit any defects in gluconeogenesis or hypoglycemia under the conditions examined. To determine the contribution of hepatic amino acid metabolism to exercise performance and metabolism, we subjected LS-*Gpt2* mice to an acute, graded intensity treadmill exercise challenge that was designed to evaluate exercise endurance. Compared to littermate WT mice, LS-*Gpt2*-/- mice ran for a similar duration and distance (Figure 1B). After exercise, fasted blood glucose and lactate concentrations were similar between genotypes during the 1-hour recovery period (Figure 1C and Figure 1D) and there was no difference in skeletal muscle glycogen content at the end of the 1-hour recovery period (Figure 1E). This indicates that loss of ALT2 in liver does not affect exercise performance or recovery in the paradigm and parameters studied.

We also determined whether suppression of mitochondrial pyruvate metabolism in the liver could affect exercise performance given that lactate/pyruvate supported gluconeogenesis by the liver is believed to be important via the Cori cycle (Figure 2A). LS-*Mpc2*-/- mice have been previously characterized and we have demonstrated that loss of MPC2 protein destabilizes the MPC complex essentially resulting in a double knockout for MPC2 and its heterodimeric partner MPC1 (16). We found that time to exhaustion in the graded treadmill exercise paradigm was not different between LS-*Mpc2*-/- and WT littermate mice (Figure 2B), though LS-*Mpc2*-/- mice tended to run for shorter times. WT and LS-*Mpc2*-/- mice had similar blood glucose concentrations during the 1-hour recovery period (Figure 2C), but the lactate AUC was 34% greater in LS-*Mpc2*-/- mice (Figure 2D) potentially reflecting impaired Cori cycling. Gastrocnemius glycogen content was similar between WT and LS-*Mpc2* -/- mice at the end of the recovery period (Figure 2E), again suggesting that glycogen replenishment from blood glucose was unimpaired by loss of MPC in the liver. Thus, loss of the MPC in liver did not affect exercise tolerance or recovery.

### Concomitant loss of liver ALT2 and MPC2 impairs endurance exercise performance

We crossed LS-*Gpt2*-/- mice with LS-*Mpc2-/-* mice to generate double knockout (DKO) mice with liver specific deletion of both ALT2 and MPC2 (32). This blocks both pathways for alanine and pyruvate entry into gluconeogenic pathways (Figure 3A). Western blotting analyses confirmed that ALT2, MPC2, and MPC1 were absent from the livers of DKO mice, but that ALT1 was still expressed (Figure 3B). To determine the hepatocyte-intrinsic effects of DKO on glucose production, hepatocytes were isolated from littermate WT and DKO mice and glucagon-stimulated glucose production in response to pyruvate or alanine was assessed. As predicted, hepatocytes from the DKO mice produced significantly less glucose when supplied with pyruvate or alanine as a gluconeogenic substrate (Figure 3C).

We next evaluated the time to exhaustion in the graded treadmill exercise paradigm and found that, compared to WT littermate controls, mice lacking both ALT2 and MPC2 in liver exhibited a significant reduction in time to exhaustion (Figure 4A). Furthermore, DKO mice exhibited significantly lower blood glucose concentrations post-exercise as indicated by a 57% reduction compared to WT controls in fasted glucose AUC during the 1-hour recovery period (Figure 4B) while lactate AUC during recovery was increased by 93% during the same time (Figure 4C). Finally, DKO mice also had significantly less glycogen in gastrocnemius muscle compared to littermate controls after 1-hour of recovery from exercise (Figure 4D). This finding could suggest that lower liver glucose production and blood glucose drives muscle to rely on its glycogen stores for glucose demands and/or that impaired gluconeogenesis and lower blood glucose concentrations were impacting the replenishment of muscle glycogen.

### Liver-specific DKO mice are hypoglycemic during exercise with impaired anaplerotic and cataplerotic metabolism

To evaluate liver metabolism during treadmill exercise, a separate cohort of mice were shipped to the Vanderbilt Mouse Metabolic Phenotyping Core for further evaluation. Mice were then surgically catheterized to allow for continued sampling of blood during exercise to directly assess gluconeogenesis by using ^2^H/^13^C metabolic flux analysis by infusing [6,6-^2^H_2_]glucose, ^2^H_2_O, and [U-^13^C]propionate as previously described (21) and as shown in Figure 5A.

After recovery, mice underwent a 30-min acute treadmill running bout at 12 m/min. The slower speed and duration was chosen so that mice of both genotypes could complete the entire exercise bout. Thirty min of exercise tended to decrease plasma insulin concentrations compared to baseline (exercise effect: p=0.07), but insulin concentrations were not affected by genotype at either time point (Figure 5B). Although DKO mice exhibited normal glucose concentrations at the start of the exercise test and after 10 min of running, they were significantly hypoglycemic compared to WT controls after 20 and 30 min of treadmill exercise (Figure 5C). Consistent with the observed hypoglycemia, quantification of nutrient fluxes revealed that endogenous glucose production, total gluconeogenesis, and gluconeogenesis from phosphoenolpyruvate (PEP) were significantly reduced in DKO mice compared to WT controls at the start of the exercise bout and at all times during exercise (Figure 5C). Interestingly, gluconeogenesis from glycerol was not affected by loss of MPC and ALT2 activity at baseline, and was actually increased in DKO mice versus WT controls after 30 min of exercise (Figure 5C). Finally, glucose production from glycogenolysis, which should not be directly impacted by loss of the MPC and ALT2, was not affected at baseline, but was markedly reduced in DKO mice compared to WT controls at all time points thereafter (Figure 5C).

In addition to glucose fluxes, mitochondrial oxidative fluxes were determined in WT and DKO mice (Figure 6A). Consistent with the lower gluconeogenesis from PEP, cataplerosis (V_PCK_) was decreased in DKO mice at rest and during exercise (Figure 6B). Anaplerosis and related fluxes (V_PC_, V_PCC_, V_LDH_) were lower at rest and during exercise in DKO mice compared to WT mice (Figure 6B). This included the flux of non-phosphoenolpyruvate anaplerotic sources to pyruvate (V_LDH_), which encompasses lactate and alanine conversion to pyruvate. TCA cycle fluxes (V_CS_ and V_SDH_) and pyruvate cycling (V_PK+ME_) increased during exercise in WT mice(Figure 6B). However, the rise in these fluxes was markedly abrogated in DKO mice. Together, these results show that inhibiting key nodes regulating pyruvate entry and generation in the mitochondria impairs the oxidative and gluconeogenic capacity of the liver and limits the ability to the liver to meet the glucose demands of working muscle.

We assessed the protein expression of the enzymes catalyzing key reactions in these pathways, including phosphoenolpyruvate carboxykinase 1 (PEPCK1), pyruvate carboxylase (PC), citrate synthase (CS), and lactate dehydrogenase (LDH) in liver samples collected from these mice at the end of the study. Although flux through these metabolic pathways was markedly suppressed, the protein abundance of these enzymes was not affected in liver of the DKO mice compared to WT controls (Figure 6C). In addition, the expression of subunits of the five electron transport chain complexes was also not affected by loss of ALT2 and the MPC, suggesting that reduced flux through these metabolic reactions is due to impaired pyruvate and alanine metabolism.

## DISCUSSION

Accelerated hepatic glucose production via glycogenolysis and gluconeogenesis during exercise plays an important role in fueling muscle contraction via the activities of the Cori and Cahill cycles. To evaluate the role of hepatic mitochondrial pyruvate and amino acid metabolism, we evaluated exercise performance and metabolic responses to acute exercise in mice lacking MPC2, ALT2, or both proteins in a hepatocyte-specific manner. We found that deletion of either MPC2 or ALT2 alone did not significantly impair exercise performance, but that DKO mice fatigued sooner and were hypoglycemic during and after the exercise challenge. Moreover, using sophisticated ^2^H/^13^C metabolic flux analyses we demonstrated that DKO mice exhibited lower endogenous glucose production at rest and during exercise (owing to a decrease in both glycogenolysis and gluconeogenesis). The decline in gluconeogenesis was linked to decreased anaplerotic, cataplerotic, and TCA cycle fluxes. These data suggest that metabolic inhibition of both the Cori and Cahill cycles is required to jointly lower hepatic glucose production and constrain exercise performance in mice.

The liver has an extraordinary ability to extract multiple types of nutrients from the blood and channel them to gluconeogenesis, which allows the liver to effectively maintain glucose homeostasis and meet the glucose demands of working muscle. The data presented herein highlight the important roles that coordinated use of different fuel sources in liver mitochondrial metabolism plays in maintaining blood glucose concentrations during exercise and how this may impact prolonged exercise performance. The present findings demonstrate that only in the context of preventing the use of multiple gluconeogenic precursors (i.e., a major interruption in the ability of the liver to transition to the gluconeogenic mode) that accelerated gluconeogenesis during exercise and exercise performance are impacted.

Previously we have shown that the modest effects of deleting the MPC in the liver on blood glucose under fasted conditions were mitigated, at least in part, by compensatory pyruvate-alanine cycling via ALT2 (16). In other work, disruption of the gene encoding ALT2 (*Gpt2*) in hepatocytes resulted in little discernable effect on glucose homeostasis in fasted lean mice and this is likely due to alanine-pyruvate cycling via the intact MPC in those mice (15). Similarly, liver-specific deletion of either *Mpc2* or *Gpt2* had no effect on blood glucose concentrations after exercise. However, we show that combined loss of *Gpt2* and *Mpc2* leads to marked impairment in hepatic gluconeogenic, cataplerotic, anaplerotic, and overall TCA flux especially during treadmill exercise. The phenotype of these mice during exercise is quite consistent with recently reported effects of deleting hepatic PEPCK1 (21), which also impairs flux of pyruvate and alanine into gluconeogenic pathways. Interestingly, we have found that DKO mice can maintain normal blood glucose during 16 h fasted conditions even with concomitant liver-specific suppression of glycerol mediated gluconeogenesis (32). This is likely due to the gluconeogenic contributions of other tissues (kidney and small intestine) under chronic fasted conditions (3, 17, 24). The inability of DKO mice to maintain normoglycemia during exercise highlights the tremendous demand for peripheral glucose supply during exercise and the potential inability for other gluconeogenic tissues to compensate in the context of an acute and robust demand for hepatic gluconeogenesis.

Results from the metabolic flux studies showed that loss of both MPC and ALT2 resulted in reduced gluconeogenesis at rest and an attenuated increase with the onset of exercise. Interestingly, gluconeogenesis from glycerol began to increase in DKO mice after 20 min of continuous aerobic activity. There are two pathways for gluconeogenesis from glycerol. The direct pathway essentially reverses some steps of glycolysis and another indirect pathway that requires mitochondrial metabolism of glycerol as pyruvate. Our recent work has demonstrated that loss of the MPC impairs glycerol metabolism through the indirect pathway, but does not affect overall glycerol-mediated gluconeogenesis, which is the predominant pathway in liver (32). However, increase in gluconeogenesis from glycerol was not sufficient to compensate for the dramatic reductions in gluconeogenesis from pyruvate/lactate and alanine, as the absolute contribution of glycerol to overall glucose production was quite low. This resulted in reduced blood glucose and elevated blood lactate concentrations during and after exercise. Reductions in blood glucose and accumulation of blood lactate beyond the rate of clearance, correlate well with exercise fatigue during fasting (29), which could explain reduced performance in DKO mice, though we did not perform experiments to determine the relative roles of each in the present study.

We found that DKO mice exhausted more quickly when a graded intensity test that plateaued at 30 m/min was employed, suggesting that liver glucose production is an important factor in an exercise paradigm with a sustained workload. This paradigm was designed to measure endurance in the context of a submaximal workload. On the other hand, despite the striking defects in gluconeogenesis, maximal running speed in an exercise stress test (analogous to a VO_2_ max test) in the DKO mice was not different than WT control mice (Supplemental Figure 1). This higher intensity protocol, likely tests aerobic capacity and our findings suggest that in this context hepatic glucose production is not limiting for exercise capacity. It is important to note that mice used for exercise experiments only engaged in acute bouts of exercise, and how inhibition of MPC, ALT2, or both affects adaptations to chronic exercise training remains unexplored. However, recent work with another model with liver-specific defects in gluconeogenesis, the liver-specific PEPCK knockout mouse, revealed no defects in adaptations to exercise training (21).

In conclusion, combined deletion of *Mpc2* and *Gpt2* in liver of mice markedly impairs hepatic gluconeogenesis and leads to hypoglycemia, hyperlactatemia, and reduced exercise performance in a graded intensity treadmill protocol. Moreover, loss of mitochondrial pyruvate and alanine metabolism results in markedly reduced flux through a variety of mitochondrial pathways in liver during exercise. These findings highlight the importance of hepatic glucose production in many exercise paradigms.

## CURRENT ADDRESSES

Michael R. Martino is currently affiliated with the Medical University of South Carolina

## AUTHOR CONTRIBUTIONS

Michael R. Martino: Conceived and designed research, performed experiments, analyzed data, interpreted results of experiments, prepared figures, drafted manuscript, edited and revised manuscript, approved final version of manuscript.

Mohammad Habibi: Performed experiments, analyzed data, interpreted results of experiments, prepared figures, drafted manuscript, edited and revised manuscript, approved final version of manuscript.

Daniel Ferguson: Performed experiments, analyzed data, interpreted results of experiments, prepared figures, drafted manuscript, edited and revised manuscript, approved final version of manuscript.

Rita T. Brookheart: Interpreted results of experiments, edited and revised manuscript, approved final version of manuscript.

John P. Thyfault: Interpreted results of experiments, edited and revised manuscript, approved final version of manuscript.

Gretchen A. Meyer: Interpreted results of experiments, edited and revised manuscript, approved final version of manuscript.

Louise Lantier: Conceived and designed research, performed experiments, analyzed data, interpreted results of experiments, prepared figures, drafted manuscript, edited and revised manuscript, approved final version of manuscript.

Curtis C. Hughey: Conceived and designed research, performed experiments, analyzed data, interpreted results of experiments, prepared figures, drafted manuscript, edited and revised manuscript, approved final version of manuscript.

Brian N. Finck: Conceived and designed research, analyzed data, interpreted results of experiments, prepared figures, drafted manuscript, edited and revised manuscript, approved final version of manuscript.

---

**Supplemental Figure 1.**
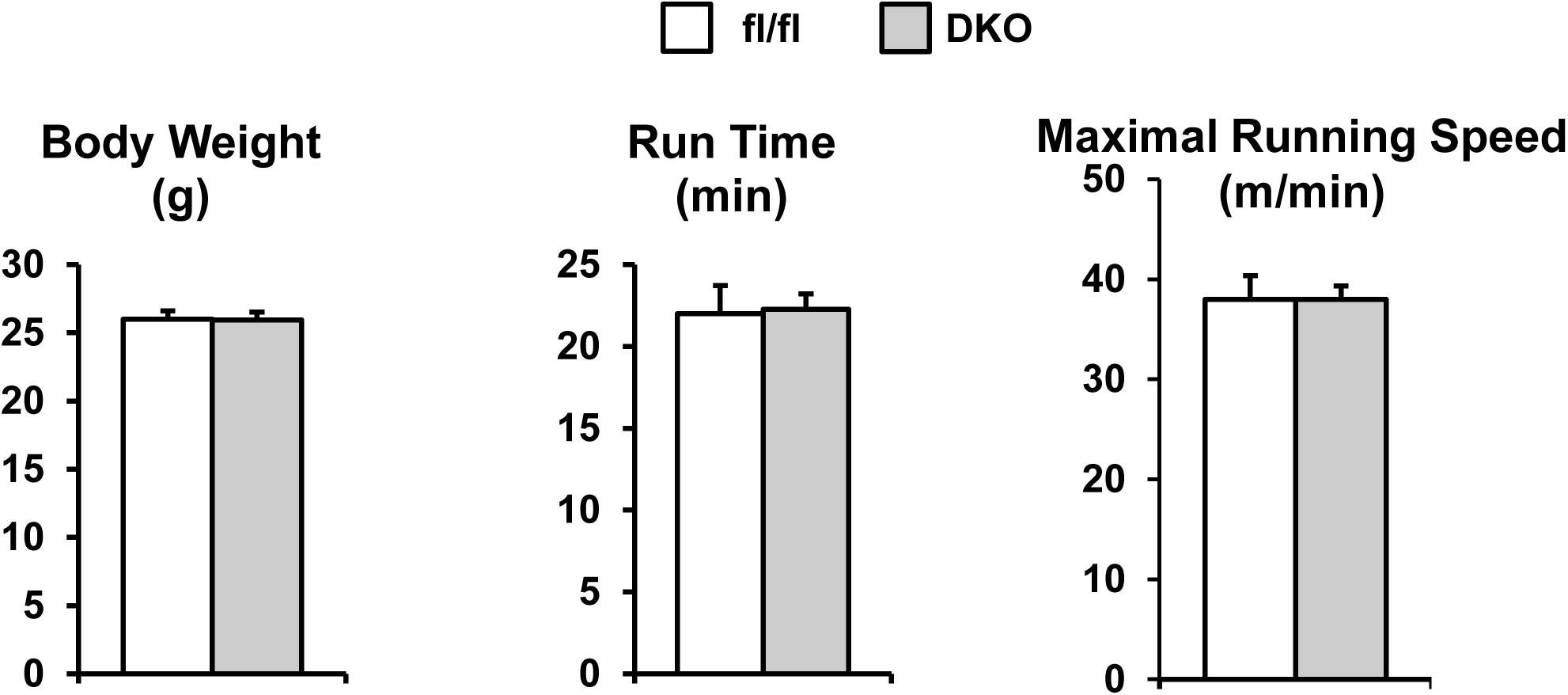
Body weight, run time, and maximal running speed of WT and DKO in an acute exercise stress test. After 4 weeks of quarantine and acclimation at Vanderbilt University, surgically-naïve mice underwent an exercise stress test to assess maximal running speed. In this paradigm, the treadmill speed starts at 10 m/min and the speed increases 4 m/min every 3 min until exhaustion.

## Literature Cited

1. Ayala JE, Bracy DP, Malabanan C, James FD, Ansari T, Fueger PT, McGuinness OP, Wasserman DH. Hyperinsulinemic-euglycemic clamps in conscious, unrestrained mice. J Vis Exp 2011.

2. Bricker DK, Taylor EB, Schell JC, Orsak T, Boutron A, Chen YC, Cox JE, Cardon CM, Van Vranken JG, Dephoure N, Redin C, Boudina S, Gygi SP, Brivet M, Thummel CS, Rutter J. A mitochondrial pyruvate carrier required for pyruvate uptake in yeast, Drosophila, and humans. Science 337: 96–100, 2012.

3. Cappel DA, Deja S, Duarte JAG, Kucejova B, Inigo M, Fletcher JA, Fu X, Berglund ED, Liu T, Elmquist JK, Hammer S, Mishra P, Browning JD, Burgess SC. Pyruvate-Carboxylase-Mediated Anaplerosis Promotes Antioxidant Capacity by Sustaining TCA Cycle and Redox Metabolism in Liver. Cell Metab 29: 1291–1305 e1298, 2019.

4. Chen Z, Vigueira PA, Chambers KT, Hall AM, Mitra MS, Qi N, McDonald WG, Colca JR, Kletzien RF, Finck BN. Insulin resistance and metabolic derangements in obese mice are ameliorated by a novel peroxisome proliferator-activated receptor gamma-sparing thiazolidinedione. J Biol Chem 287: 23537–23548, 2012.

5. Donovan CM and Sumida KD. Training improves glucose homeostasis in rats during exercise via glucose production. The American journal of physiology 258: R770–776, 1990.

6. Felig P, Owen OE, Wahren J, Cahill GF, Jr. Amino acid metabolism during prolonged starvation. The Journal of clinical investigation 48: 584–594, 1969.

7. Felig P, Pozefsk T, Marlis E, Cahill GF. Alanine: Key Role in Gluconeogenesis. Science 167: 1003–1004, 1970.

8. Felig P and Wahren J. Amino acid metabolism in exercising man. The Journal of clinical investigation 50: 2703–2714, 1971.

9. Gray LR, Sultana MR, Rauckhorst AJ, Oonthonpan L, Tompkins SC, Sharma A, Fu X, Miao R, Pewa AD, Brown KS, Lane EE, Dohlman A, Zepeda-Orozco D, Xie J, Rutter J, Norris AW, Cox JE, Burgess SC, Potthoff MJ, Taylor EB. Hepatic Mitochondrial Pyruvate Carrier 1 Is Required for Efficient Regulation of Gluconeogenesis and Whole-Body Glucose Homeostasis. Cell Metab 22: 669–681, 2015.

10. Gray LR, Tompkins SC and Taylor EB. Regulation of pyruvate metabolism and human disease. Cellular and Molecular Life Sciences 71: 2577–2604, 2014.

11. Herzig S, Raemy E, Montessuit S, Veuthey JL, Zamboni N, Westermann B, Kunji ER, Martinou JC. Identification and functional expression of the mitochondrial pyruvate carrier. Science 337: 93–96, 2012.

12. Hodges WT, Jarasvaraparn C, Ferguson D, Griffett K, Gill LE, Chen Y, Ilagan MXG, Hegazy L, Elgendy B, Cho K, Patti GJ, McCommis KS, Finck BN. Mitochondrial pyruvate carrier inhibitors improve metabolic parameters in diet-induced obese mice. J Biol Chem 298: 101554, 2022.

13. Hughey CC, James FD, Bracy DP, Donahue EP, Young JD, Viollet B, Foretz M, Wasserman DH. Loss of hepatic AMP-activated protein kinase impedes the rate of glycogenolysis but not gluconeogenic fluxes in exercising mice. J Biol Chem 292: 20125–20140, 2017.

14. Lindblom P, Rafter I, Copley C, Andersson U, Hedberg JJ, Berg A-L, Samuelsson A, Hellmold H, Cotgreave I, Glinghammar B. Isoforms of alanine aminotransferases in human tissues and serum—differential tissue expression using novel antibodies. Archives of biochemistry and biophysics 466: 66–77, 2007.

15. Martino MR, Gutierrez-Aguilar M, Yiew NKH, Lutkewitte AJ, Singer JM, McCommis KS, Ferguson D, Liss KHH, Yoshino J, Renkemeyer MK, Smith GI, Cho K, Fletcher JA, Klein S, Patti GJ, Burgess SC, Finck BN. Silencing alanine transaminase 2 in diabetic liver attenuates hyperglycemia by reducing gluconeogenesis from amino acids. Cell Rep 39: 110733, 2022.

16. McCommis KS, Chen Z, Fu X, McDonald WG, Colca JR, Kletzien RF, Burgess SC, Finck BN. Loss of Mitochondrial Pyruvate Carrier 2 in the Liver Leads to Defects in Gluconeogenesis and Compensation via Pyruvate-Alanine Cycling. Cell Metab 22: 682–694, 2015.

17. Mutel E, Gautier-Stein A, Abdul-Wahed A, Amigó-Correig M, Zitoun C, Stefanutti A, Houberdon I, Tourette JA, Mithieux G, Rajas F. Control of blood glucose in the absence of hepatic glucose production during prolonged fasting in mice: induction of renal and intestinal gluconeogenesis by glucagon. Diabetes 60: 3121–3131, 2011.

18. Okun JG, Rusu PM, Chan AY, Wu Y, Yap YW, Sharkie T, Schumacher J, Schmidt KV, Roberts-Thomson KM, Russell RD, Zota A, Hille S, Jungmann A, Maggi L, Lee Y, Blüher M, Herzig S, Keske MA, Heikenwalder M, Müller OJ, Rose AJ. Liver alanine catabolism promotes skeletal muscle atrophy and hyperglycaemia in type 2 diabetes. Nature Metabolism 3: 394–409, 2021.

19. Ouyang Q, Nakayama T, Baytas O, Davidson SM, Yang C, Schmidt M, Lizarraga SB, Mishra S, EI-Quessny M, Niaz S, Gul Butt M, Imran Murtaza S, Javed A, Chaudhry HR, Vaughan DJ, Hill RS, Partlow JN, Yoo S-Y, Lam A-TN, Nasir R, Al-Saffar M, Barkovich AJ, Schwede M, Nagpal S, Rajab A, DeBerardinis RJ, Housman DE, Mochida GH, Morrow EM. Mutations in mitochondrial enzyme GPT2 cause metabolic dysfunction and neurological disease with developmental and progressive features. Proceedings of the National Academy of Sciences 113: E5598–E5607, 2016.

20. Radziuk J and Pye S. Hepatic glucose uptake, gluconeogenesis and the regulation of glycogen synthesis. Diabetes/metabolism research and reviews 17: 250–272, 2001.

21. Rome FI, Shobert GL, Voigt WC, Stagg DB, Puchalska P, Burgess SC, Crawford PA, Hughey CC. Loss of hepatic phosphoenolpyruvate carboxykinase 1 dysregulates metabolic responses to acute exercise but enhances adaptations to exercise training in mice. Am J Physiol Endocrinol Metab 324: E9–E23, 2023.

22. Rubin RP. Carl and Gerty Cori: A collaboration that changed the face of biochemistry. J Med Biogr 29: 143–148, 2021.

23. Ruderman NB. Muscle amino acid metabolism and gluconeogenesis. Annual review of medicine 26: 245–258, 1975.

24. She P, Burgess SC, Shiota M, Flakoll P, Donahue EP, Malloy CR, Sherry AD, Magnuson MA. Mechanisms by which liver-specific PEPCK knockout mice preserve euglycemia during starvation. Diabetes 52: 1649–1654, 2003.

25. Snell K. Muscle alanine synthesis and hepatic gluconeogenesis. Biochemical Society transactions 8: 205–213, 1980.

26. Snell K and Duff DA. Alanine and glutamine formation by muscle. Biochemical Society transactions 8: 501–504, 1980.

27. Sumida K and Donovan CM. Enhanced hepatic gluconeogenic capacity for selected precursors after endurance training. Journal of Applied Physiology 79: 1883–1888, 1995.

28. Suzuki Y, Lanner C, Kim JH, Vilardo PG, Zhang H, Yang J, Cooper LD, Steele M, Kennedy A, Bock CB, Scrimgeour A, Lawrence JC, Jr., DePaoli-Roach AA. Insulin control of glycogen metabolism in knockout mice lacking the muscle-specific protein phosphatase PP1G/RGL. Mol Cell Biol 21: 2683–2694, 2001.

29. Travassos PB, Godoy G, De Souza HM, Curi R, Bazotte RB. Performance during a strenuous swimming session is associated with high blood lactate: pyruvate ratio and hypoglycemia in fasted rats. Braz J Med Biol Res 51: e7057, 2018.

30. Wasserman DH. Four grams of glucose. American Journal of Physiology - Endocrinology And Metabolism 296: E11–E21, 2009.

31. Wasserman DH, Williams PE, Lacy DB, Green DR, Cherrington AD. Importance of intrahepatic mechanisms to gluconeogenesis from alanine during exercise and recovery. Am J Physiol 254: E518–525, 1988.

32. Yiew NKH, Ferguson D, Cho K, Deja S, Jarasvaraparn C, Lutkewitte AJ, Mukherjee S, Fu X, Singer JM, Patti GJ, Burgess SC, Finck BN. Effects of hepatic mitochondrial pyruvate carrier deficiency on de novo lipogenesis and glycerol-mediated gluconeogenesis in mice. bioRxiv 2023.2002.2017.528992, 2023.

